# Polyphosphate Discriminates Protein Conformational Ensembles More Efficiently than DNA Promoting Diverse Assembly and Maturation Behaviors

**DOI:** 10.1101/2024.10.21.619443

**Authors:** Saloni Goyal, Divya Rajendran, Anup Kumar Mani, Athi N. Naganathan

## Abstract

Disordered proteins and domains often assemble into condensates with polyanionic nucleic acids, primarily via charge complementarity, regulating numerous cellular functions. However, the assembly mechanisms associated with the other abundant and ubiquitous, anionic, stress-response regulating polymer, polyphosphate (polyP), is less understood. Here, we employ the intrinsically disordered DNA binding domain (DBD) of cytidine repressor (CytR) from *E.coli* to study the nature of assembly processes with polyP and DNA. CytR forms metastable liquid-like condensates with polyP and DNA, while undergoing liquid-to-solid transition in the former and dissolving in the latter. On mutationally engineering the ensemble to exhibit more or less structure and dimensions than the WT, the assembly process with polyP is directed to either condensates with partial time-dependent dissolution or spontaneous aggregation, respectively. On the other hand, the CytR variants form *only* liquid-like but metastable droplets with DNA which dissolve within a few hours. Polyphosphate induces large secondary-structure changes, with two of the mutants adopting polyproline II-like structures within droplets, while DNA has only minimal structural effects. Our findings reveal how polyphosphate can more efficiently discern conformational heterogeneity in the starting protein ensemble, its structure, and compactness, with broad implications in assembly mechanisms involving polyP and stress response in bacterial systems.

## Introduction

Biomolecular condensates are mesoscale membrane-less structures that facilitate various cellular processes from stress response and signal transduction to RNA processing and chromatin organization.^1–5^ Physically, condensates are formed by the demixing or phase separation of bio-macromolecules (i.e. proteins, DNA, RNA) into a polymer-rich or a condensed phase and a solvent-rich phase. The phase separation can be driven by homotypic or heterotypic interactions depending on the nature of demixing species, with a host of factors including the concentration and composition of constituent species, specific ions, ionic strength, temperature, and pH governing the transition.^6–8^ Both folded and disordered regions of proteins can form condensates, with a lot more propensity and prevalence noted in the latter. Biomolecular condensates are also implicated in numerous diseases, and specifically in neurodegenerative disorders, with the condensates or sometimes the dilute phase proposed to seed the formation of aggregates and amyloids.^5,9–12^

Complex coacervation, wherein oppositely charged macromolecules condense and promote phase transition, is often observed in DNA- and RNA-binding domains that harbor excess positive charges to bind the anionic counterpart.^7,13–19^ However, the role of inorganic polyphosphates (polyP), a series of phosphate units (Pi) linked together via phosphoanhydride bonds (Figure 1A), in condensate formation and maturation has been less explored. PolyP is a ubiquitous molecule that is produced under stress conditions like oxidative stress, heat shock, nutrition limitation, etc., in all organisms.^20–22^ In fact, the polymer length of polyP and the concentration vary among different species ranging from few tens to thousands, reaching up to 50 mM in bacterial cells under stress conditions.^20^ Apart from stress response, it has been implicated in numerous processes ranging from modulation of nucleoid structure to biofilm formation.^23–28^ Intriguingly, polyP is able to act both as a chaperone^29^ (assisting folding of proteins, preventing protein misfolding and inhibiting aggregation) and in promoting aggregation,^30,31^ indicative of more complex underlying mechanisms^32,33^ which could be protein-specific.

**Figure 1.**
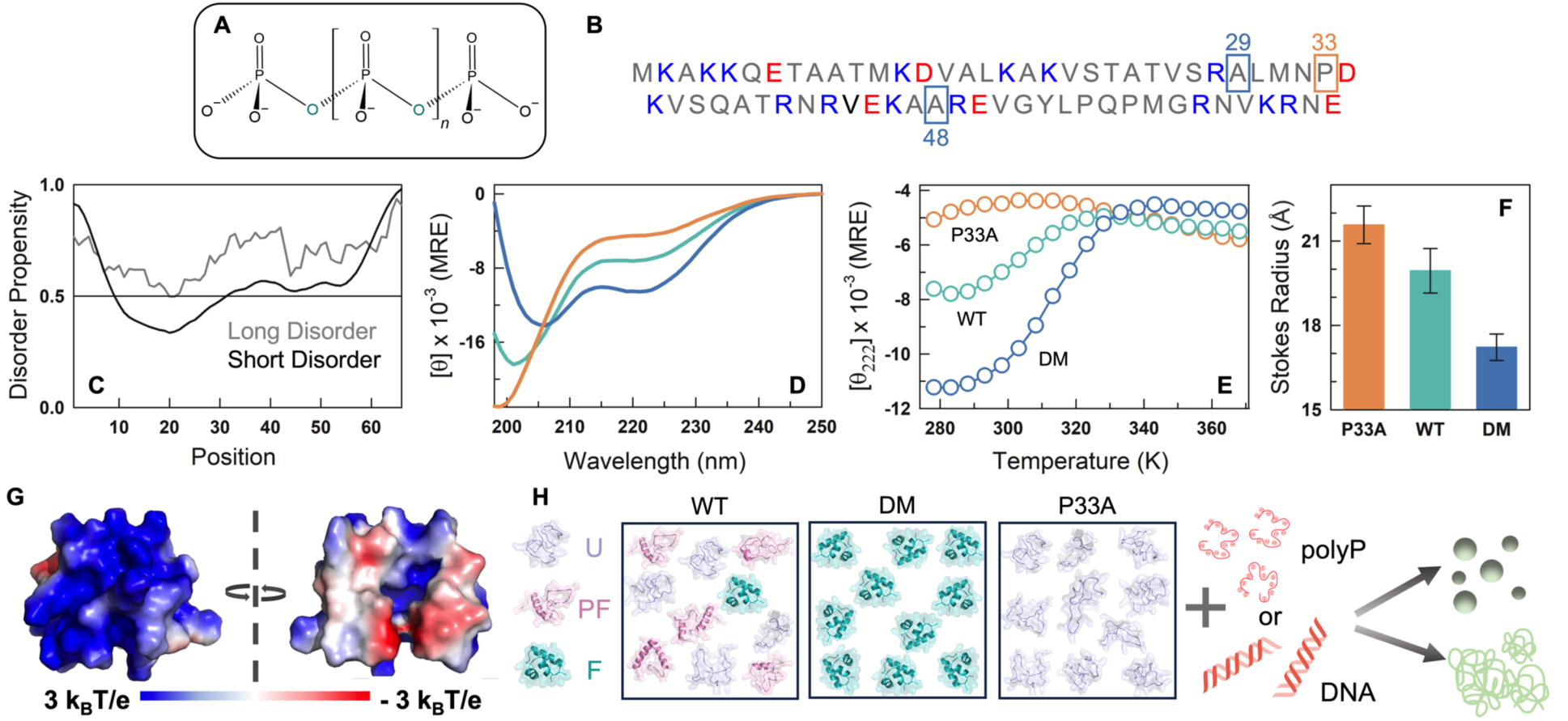
Conformational features of CytR WT and mutants. (A) Molecular structure of polyP. (B) Amino acid sequence of CytR. Blue and red boxes indicate the positions of alanine and proline mutations in DM and P33A variants, respectively. (C) Disordered propensity of CytR predicted using IUPred3^34^ with gray and black curves showing long and short disorder predictions, respectively. (D) Far-UV CD spectra of WT (green), DM (blue), and P33A (red) at 298 K in mean residue ellipticity (MRE) units of deg. cm^2^ dmol^−1^. (F) Thermal denaturation curves of CytR variants from far-UV CD experiments monitored at 222 nm and reported in MRE units. (F) Stokes Radius following the color code in panel E. (G) Electrostatic potential map^35^ of CytR in its folded conformation displaying a large positive electrostatic potential. (H) The hypothesis tested in the current work. WT (left), DM (middle), and P33A (right) ensembles could potentially form condensates or aggregates in the presence of polyP or DNA, and which could also display differential time-dependent properties. U, PF, and F represent unfolded, partially folded and folded conformations, respectively.

Based on the observations above, we pose this question: is polyphosphate more sensitive to the starting conformational ensemble than DNA leading to differential behaviors of promoting aggregation versus acting as a chaperone? If so, and given the poly-anionic nature, does it lead to aggregates or condensates or both? To answer these questions, one needs a system that is also a DBD to enable direct comparison. Second, the DBD should be mutationally tunable to populate native ensembles with different structure, compactness and helicity. Third, the charge composition and their sequence patterning should not be changed across the mutants, as this will naturally affect the binding to polyP. Here, we employ Cytidine repressor DBD (CytR DBD or CytR or CytR wildtype), an intrinsically disordered domain (Figure 1B, 1C) that adopts a folded conformation in the presence of DNA,^36^ as a model system to answer these questions. Despite its disordered nature, mutations of different types and at different positions can be introduced into CytR to make it adopt a continuum of conformations, from highly disordered to globally ordered (i.e. with a specific three-dimensional structure).^37–39^ Specifically, the WT populates a minor excited state which is fully folded (∼8-10%), but otherwise the native ensemble is disordered, with marginal helicity and a Stokes radius of 19.5 Å (green in Figure 1D, 1F).^37,40^ One of the mutants we consider is the double mutant of CytR (A29V/A48M; DM) which displays the properties of a fully folded compact domain driven by the enhanced hydrophobicity of the side-chains at positions 29 and 48; these two mutations act synergistically to switch the ensemble from being disordered to fully ordered as shown earlier.^38^ The DM therefore displays a higher helicity and lower Stokes radius of 17 Å, compared to the WT (blue in Figure 1D, 1F). The other mutant is P33A in which the minor folded state population is eliminated because of the enhanced flexibility of alanine in the place of proline.^37^ The P33A mutant is nearly fully disordered as evidenced by the Stokes radius of 21.5 Å, and minimum helicity (red in Figure 1D, 1F).

In addition, because CytR has an excess charge of +9 and a large positive electrostatic potential in its folded conformation (Figure 1G), it binds non-specifically to DNA.^41,42^ We therefore hypothesize that CytR should bind to any negatively charged polymer driven purely by electrostatic complementarity. In this work, we employ CytR WT and two mutants of CytR that exhibit diametrically opposing conformational preferences but identical charge composition and two anionic polymers (polyP and DNA), to understand the extent to which the starting ensemble (and hence the density of charges and stability or structure) dictates the nature of macromolecular assemblies and their maturation properties (Figure 1H). We find that the starting conformational ensemble is uniquely recognized by polyphosphate leading to diverse outcomes including metastable condensates, aggregates and time-dependent dissolution.

## Methods

### Purification of CytR DBD and mutants

The gene corresponding to the DNA binding domain of CytR(MKAKKQETAATMKDVALKAKVSTATVSRALMNPDKVSQATRNRVEKAAREVGYLPQPMGRNVKRNE) in the pTXB1 vector was transformed in to *E.coli* BL21 (DE3) cells and purified as described before.^41^ Identical protocols were employed for purification of the DM and P33A variants. The Stokes radius was estimated as detailed in an earlier work.^39^

### Time-dependent scattering (OD) and Thioflavin T (ThT) fluorescence measurements

Different concentrations of CytR WT were prepared by dissolving the lyophilized protein in 20 mM sodium phosphate, pH 7 (43 mM ionic strength) buffer. To generate the phase diagram, the scattering intensity (optical density) was recorded at 350 nm at protein concentrations of 5, 25, 55, 90, and 120 µM, and at different polyP concentrations of 3, 6, 11, 22, 41, and 100 µM. The measurements were acquired for 24 hours at 7-minute intervals in a microplate reader (Agilent BioTek Synergy H1) at 298 K. The sample volume in each well was 200 *μ*l. Further studies employed 90 µM protein (WT and mutants) and 22 µM polyP, unless mentioned otherwise. For the time-dependent studies recording both scattering and fluorescence, a 5 µM ThT (Sigma-Aldrich) was also added to the protein and polyP mixture; the wells were also excited at 442 nm, and the fluorescence intensity recorded at an emission wavelength of 485 nm. Ionic strength dependent experiments were carried out at 20 mM sodium phosphate buffer, pH 7.0 (43 mM effective ionic strength) with the addition of NaCl for a final ionic strength values spanning 60 - 200 mM. The OD values are reported directly from the plate reader without correction for pathlength which is 5 mm in this experiment.

For the scattering measurements in the presence of DNA, a range of CytR WT concentrations (2 – 100 µM) were used, and measurements were acquired with 1 µM of 45-bp dsDNA (5’ TGGTGGGTAAATTTATGCAACGCATTTGCGTCATGGTGATGAGTA 3’). Further studies with DNA were done at 25 µM protein (WT and mutants) concentration.

### NHS-rhodamine labeling

NHS-rhodamine (ThermoFisher Scientific, USA) was dissolved in DMSO, and protein was dissolved in conjugation buffer at pH 7.4 (as per manufacturer’s protocol). The protein solution was added to 5X – 7X molar excess of dye and incubated at 25 °C for 2 hours at mild shaking conditions in dark. The excess dye was removed by desalting using a 26/10 HiPrep desalting column (Cytiva). The labeled protein was eluted in 20 mM sodium phosphate buffer at pH 7.

### Flow chamber preparation for imaging experiments

Imaging experiments were done using 22×22 mm coverslips and glass slides (Blue Star, India). Coverslips and glass slides were kept in Piranha solution (3:1 v/v sulphuric acid and hydrogen peroxide) for 30 minutes and then washed thoroughly with MilliQ water. The slides and coverslips were then air-dried and wiped with absolute ethanol. Clean slides and coverslips were used to prepare flow chambers to observe the condensate formation. To prepare the flow chamber, the double-stick tape was sandwiched between the glass slide and coverslip. The chamber was then sealed from three sides to avoid sample leakage.

### Fluorescence microscopy

The condensates were observed under 100X/1.45 NA oil immersion objective at room temperature using Olympus IX83 inverted fluorescence microscope. 10% of the labeled protein was mixed with unlabeled protein, and the sample was loaded into the flow chamber immediately after adding polyP/DNA. An anti-fading cocktail containing 20 µg/ml glucose oxidase, 1.4 µg/ml catalase, 20 mM glucose and 10 mM dithiothreitol was used to prevent photobleaching of the sample.^43^ The sample was imaged at different time points under DIC and fluorescence mode using an appropriate fluorescence channel at 2048 x 2048 pixels (images for WT in the presence of polyP were acquired at 1024 × 1024 pixels) with 8-bit depth resolution. Images were analyzed using cellSens Dimensions Desktop (provided with the instrument) and ImageJ.^44^

### Fluorescence recovery after photobleaching (FRAP)

For FRAP experiments, 10% and 5% of labeled protein were used for polyP and DNA studies, respectively. Anti-fading cocktail was added to the protein sample to prevent the passive photobleaching. The samples were loaded into the flow chamber at different time points (0, 30, and 60 min), and the experiment was performed in an Olympus Fluoview FV3000 confocal microscope with a 100X/1.45 NA oil immersion objective. A 561 nm laser was used to visualize and bleach (with 100% laser power) the condensates at the center. Fluorescence intensity was acquired for five different regions of interest (ROIs) with the same diameter; ROI-1 for the actual bleaching region, ROI-2 for the neighboring condensate for passive bleaching correction and ROI-3,4 and 5 outside the condensates to record background intensity for background correction. The images were acquired at 1024 × 1024 pixels with 12-bit resolution. Data was processed using cellSens Dimensions Desktop software and MATLAB. The raw data was corrected using the following equation^45^

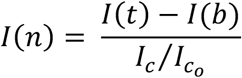

where, *I*(*t*) is the fluorescence intensity at time t, *I*(*b*) is the average of background intensity for ROI-3,4, and 5, *I_c_* is the fluorescence intensity of ROI-2 after photobleaching and 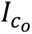 is the fluorescence intensity before photobleaching. The data was normalized (I) using:

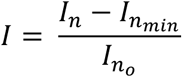

where *I_n_* is corrected fluorescence intensity at time t and 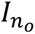 is the intensity before bleaching and 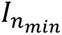 is the minimum intensity. The average of corrected and normalized fluorescence intensity from different experiments was taken, and the data was plotted as a function of time.

### Circular dichroism (CD)

Far-UV CD spectra were recorded at protein concentrations 55 µM and 25 µM immediately after the addition of 22 µM polyP and 1 µM DNA, respectively. The spectra were acquired as a function of time for 4 hours at an interval of 15 minutes in the wavelength range of 250-200 nm at 298 K with a scanning speed of 10 nm/min in a Jasco J-815 spectropolarimeter. The mean helical content for the different CytR variants is estimated by taking the ratio between the signal at 222 nm with that expected of a 100% helical protein (−40,000 MRE units of deg. cm^2^ dmol^−1^).

### Fourier-transform infrared (FTIR) spectroscopy

FTIR scans were performed using a Perkin Elmer Spectrum Two Spectrometer equipped with a DTGS detector in the attenuated total reflectance (ATR) mode. 5 µl of protein sample was added to the diamond crystal, and the spectrum was acquired in the range of 4000-400 cm^−1^ using an average of 50 scans at a resolution of 4 cm^−1^. Baseline correction was done with MilliQ water to minimize the interference due to H_2_O before spectral acquisition.

## Results and Discussion

### CytR WT and DM form condensates with polyphosphate while P33A aggregates

Inorganic polyP_45_, i.e. polyphosphate with 45 Pi units and with a net-charge of −47, is employed in our studies and is referred to as polyP in the text below. We explored a range of solution conditions involving two-component mixtures of WT and polyP ([polyP] of 3 - 100 *μ*M and [WT] of 5 – 125 *μ*M at 20 mM phosphate buffer, pH 7 and 298 K) to map the phase diagram (Figure 2A). Solutions containing [WT]>25 *μ*M and [polyP]>10 *μ*M consistently showed higher turbidity (turbidity at 350 nm). We therefore chose [polyP] of 22 *μ*M for further experiments. DIC (differential interference contrast) microscopy reveals droplet-shaped condensates with a diameter of ∼1-4 microns on average (Figure 2B; also *vide infra*), which is also confirmed by fluorescence microscopy of NHS-rhodamine-labeled protein (Figure 2C). The number of droplets increase with increasing [WT] in the range between 25 – 90 *μ*M consistent with OD measurements (Figure 2D). These observations establish that the WT CytR undergoes phase-separation into condensates with polyP, and that turbidity can be employed as a reasonable proxy for phase separation in this system.

**Figure 2.**
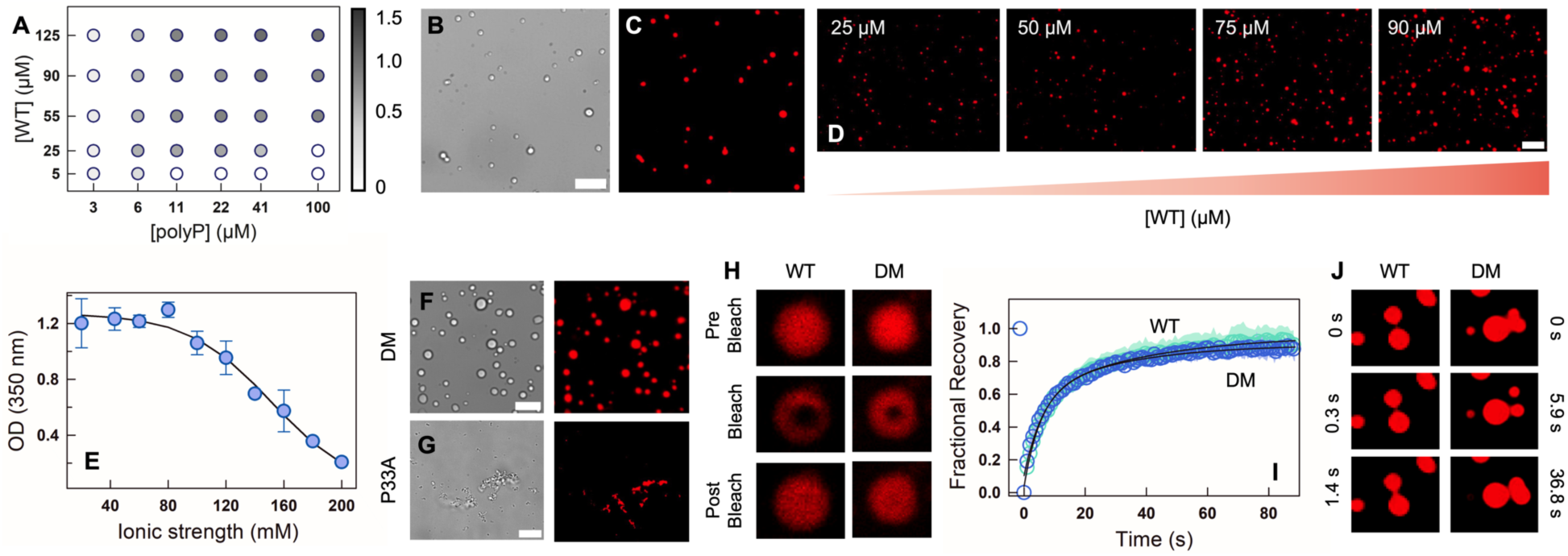
CytR undergoes phase separation with polyP *in vitro*. (A) Phase diagram illustrating the phase separation of WT at different protein and polyP concentrations. Empty circles represent no phase separation, and filled circles represent the extent of phase separation following the color bar which indicates the OD (turbidity) at 350 nm. (B, C) Representative DIC (B) and fluorescence (C) microscopy images of WT showing condensate formation in the presence of polyP. (D) Fluorescence microscopy images of NHS-rhodamine labeled WT at different protein concentrations in the presence of 22 µM PolyP. The scale bar is 10 µm. (E) Ionic strength dependence of turbidity at fixed WT (90 µM) and polyP (22 µM) concentrations. The error bar indicates the spread from experimental replicates (N=2). (F, G) DIC (left) and Fluorescence (right) microscopy images of NHS-rhodamine labeled DM (F) and P33A (G) in the presence of polyP. The scale bar is 10 µm. (H) Representative fluorescence microscopy images of condensates during FRAP on WT (left) and DM (right) immediately after polyP addition. (I) FRAP recovery curves at 0 h for DM (blue) and WT (green). The data represents an average of five experiments (N=5) and the errors (shaded areas) are smaller than the size of the circles. (J) Time-lapse fluorescence images showing a dripping event for the WT (left column) and fission-fusion events for DM (right column).

Given that CytR carries a large excess positive charge (+9) and polyP is an anionic polymer with no structural preference (unlike DNA), the interaction and the resultant assembly process is expected to be driven by non-specific electrostatic interactions. To test this, we use a [WT]:[polyP] of 90:22 *μ*M and measure OD as a function of ionic strength at pH 7. The OD is observed to be independent of ionic strength till ∼80 mM following which it linearly decreases with near-zero turbidity at 200 mM (Figure 2E). Under physiological ionic strength conditions of 150-160 mM, the OD value is still ∼0.6 and with droplets evident in fluorescence microscopy (Figure S1).

The double mutant (DM) which is folded and compact, also undergoes spontaneous phase separation at the same concentrations as the WT when mixed with polyP, while the P33A mutant aggregates on addition of polyP (Figure 2F, 2G). The fluorescence of the NHS-rhodamine-labeled WT and DM recover on photobleaching highlighting the liquid-like nature of the condensates (Figure 2H). Specifically, the WT displays a maximal recovery of ∼92% at 90 seconds (recovery half-time, *t*_1/2_ = 6.0 ± 2.5 seconds), and the DM recovers to ∼87% (*t*_1/2_ = 5.2 ± 1.3 seconds) (Figure 2I). We also observe multiple coalescence and de-coalescence events in both the proteins at the earliest times (Figure 2J, Supporting Movies S1 and S2).

Taken together, the fully unfolded variant of CytR (P33A) forms aggregates and the variants with more ‘folded-like’ states undergo phase separation with polyP. These results indicate that charge-composition and patterning alone do not determine the phase separation tendencies in this system. They underscore the importance of the starting ensemble, and potentially the population of partially structured substates, in determining the nature of assembly process observed.

### Maturation of polyphosphate-induced assemblies

We carried out time dependent studies in the presence of thioflavin T (ThT) enabling simultaneous measurement of OD and fluorescence in the same sample. The OD of WT-polyP mixtures show a strong time dependence with the OD decreasing from 1.2 to less than 0.9 in the first 10 minutes, but with little increase in ThT fluorescence (left panel in Figure 3A). However, the ThT fluorescence starts to increase in a sigmoidal fashion from 20 minutes. The OD, on the other hand, drops to a value of 0.75 at ∼30 minutes following which it increases and stabilizes at a value of ∼1. The WT-polyP mixture therefore forms condensates and matures into aggregates within 3 hours which is directly observable in microscopy images (Figure 3B), mirroring reports in numerous other systems that undergo droplet formation followed by a liquid-to-solid phase transition.^46–50^ Control experiments with protein alone did not reveal any changes in turbidity or aggregation in the entire 24-hour duration (Figure S2), pointing to a process entirely driven by polyP-protein interactions.

**Figure 3.**
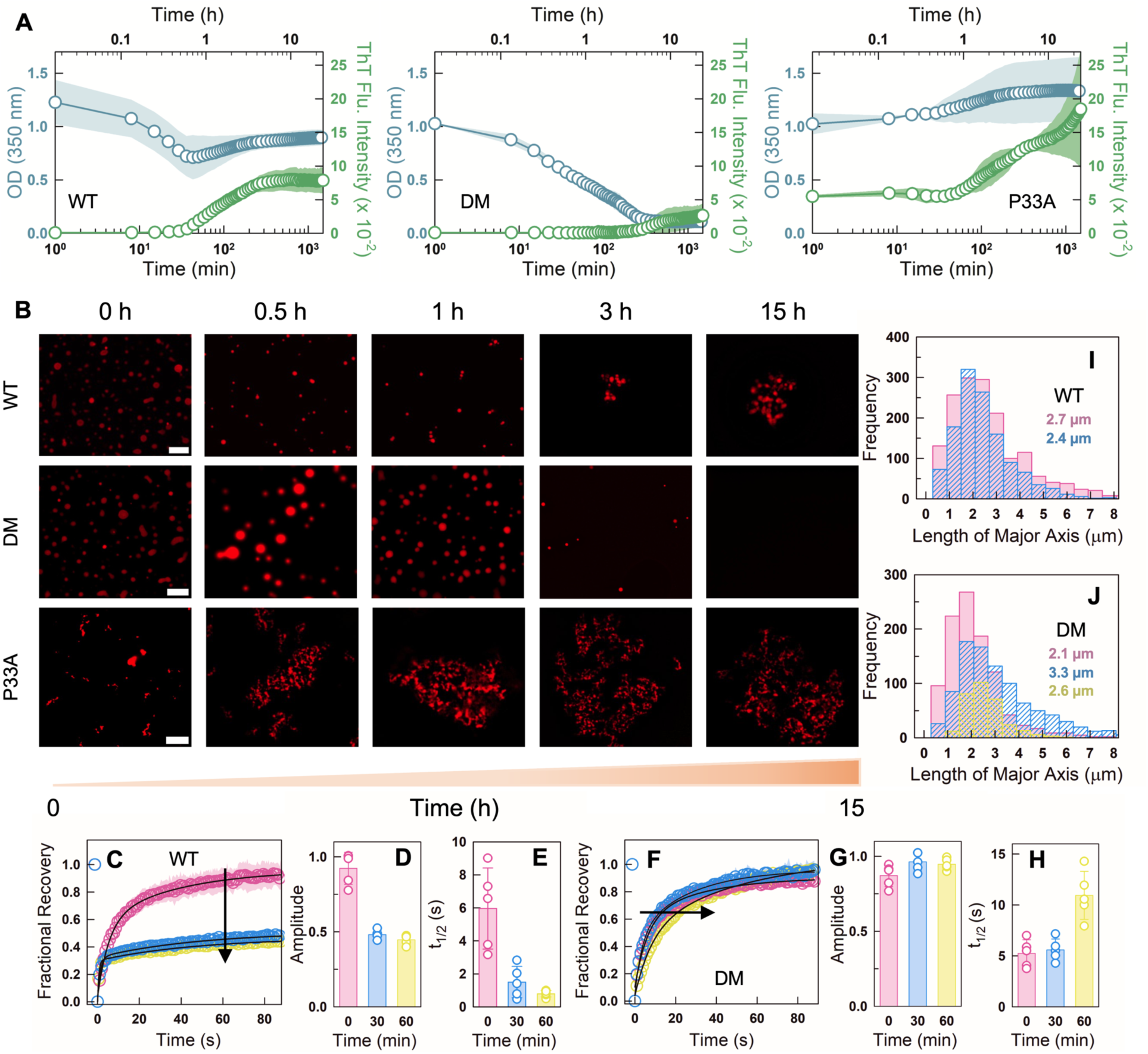
Time-dependence of polyP-induced assemblies. (A) Turbidity (blue circles and left y-axis) and Thioflavin T fluorescence intensity (green circles and right y-axis) time dependence for 90 µM WT (left panel), DM (middle panel), and P33A (right panel) in the presence of 22 µM polyP. The average from N = 2 experiments (circles) and the spread in the respective data (shaded region) are shown. (B) Representative fluorescence images of NHS-rhodamine labeled WT (top panel), DM (middle panel) and P33A (bottom panel) in the presence of polyP at different time points. The scale bar is 10 µm. (C-H) FRAP experiments for WT (panels C-E) and DM (panels F-H) in the presence of polyP at different time points - 0 min (pink), 30 min (blue), and 60 min (yellow) – and from five droplets (N = 5) and plotted as mean ± s.d. (C, F) The FRAP recovery curves of NHS-rhodamine labeled WT (panel C) and DM (panel F). The experimental errors (shaded areas) are also shown. (D, G) Recovery amplitudes or extents from FRAP experiments on the WT (panel D) and DM (panel G) at different time points. WT shows less recovery with time, indicating a liquid-to-solid transition, while the DM recovers fully. (E, H) FRAP recovery half-times for the WT (panel E) and DM (panel H) at the indicated time points. (I, J) Droplet size-distribution of WT (panel I) and DM (panel J) at different time points following the same color code in panels C-H. The numbers within the plot represent the mean droplet dimensions at the corresponding time points.

The OD of the DM-polyP mixture exhibits a similar decrease in OD starting from a value of ∼1, but this trend extends beyond 3 hours, with the OD dropping to a value of ∼0.1 at the longest time points (middle panel in Figure 3A). We do observe a marginal increase in ThT fluorescence after 2 hours, but the increase is not as significant as the WT. These observations point to a scenario wherein the DM forms condensates to start with, but which slowly dissolve with time (Figure 3B). On the other hand, the P33A mutant forms only aggregates as evidenced by the high OD and high ThT fluorescence across all time-points in the 24-hour observation window (right panel in Figure 3A, 3B). FRAP experiments follow the trends reported in OD-fluorescence measurements. In the WT, the recovery amplitude decreases from 0.92±0.11 at the zeroth minute to just 0.43±0.04 at 60 minutes (Figure 3C, 3D). The FRAP *t*_1/2_, however, drops to shorter times at *t* = 60 minutes indicating that there is small sub-set of dynamic molecules (i.e. a small percentage of mobile fraction) contributing to the rapid recovery (Figure 3E). Alternately, the DM-polyP condensates recover near fully even at the first hour, with the recovery time slowing time by a factor of 2 (Figure 3F-3H).

We further quantified the dimensions of the droplets from the microscopy images at three different time-points: 0, 30 and 60 minutes. At *t* = 0 and for the WT-polyP condensates, the mean droplet dimensions are of the order of 2.7 microns, but with a heavy right tail spanning nearly 8 microns. This tail is diminished at *t* = 30 minutes, reducing the overall dimensions of droplets (∼2.4 microns) and their number (smaller area under the curve in Figure 3I). The DM displays a behavior which is the exact opposite (Figure 3J): at *t* = 0 the mean dimensions of the condensates are 2.1 microns with no heavy right tail; larger droplets appear at *t* = 30 minutes contributing to the appearance of the heavy right tail and with the mean droplet size increasing to 3.3 microns. At longer times (*t* = 60 minutes), the number of droplets reduce and so do their mean dimensions as a result of dissolution.

### Structural changes within the condensates span over three hours

The differential behavior of the three variants raise questions on the nature of structural changes upon condensate formation and maturation. The WT CD spectrum in the dilute phase (i.e. in the absence of polyP) is characterized by minima at 202 and 222 nm with ∼17% of residual helical content (Figure 1D). Upon formation of condensates, the spectral shape shifts substantially to one with a minimum at 230 nm and with no spectral features at wavelength <215 nm (Figure 4A), demonstrating that the WT adopts a non-helical and potentially an unfolded conformation in the condensed phase at initial times. The spectrum of DM is more representative of a folded protein with minima at both 208 and 222 nm (Figure 1D), and with the signal at 222 nm indicative of a protein with nearly 30% helical content. On mixing with polyP, the spectral shift resembles that of the WT (Figure 4B), again indicative of a structural change in the condensed phase. The P33A mutant which is more unfolded (helicity ∼10%), however, exhibits a more intense spectrum with a minimum 225 nm which is representative of *β*-sheet like structures (Figure 4C; see below). These differences are more evident when the signal at 222 nm are plotted as a function of time: the WT and P33A mutant exhibit a trend quite different from that of the DM, with the latter acquiring structure with time despite unfolding on adding polyP (Figure 4D).

**Figure 4.**
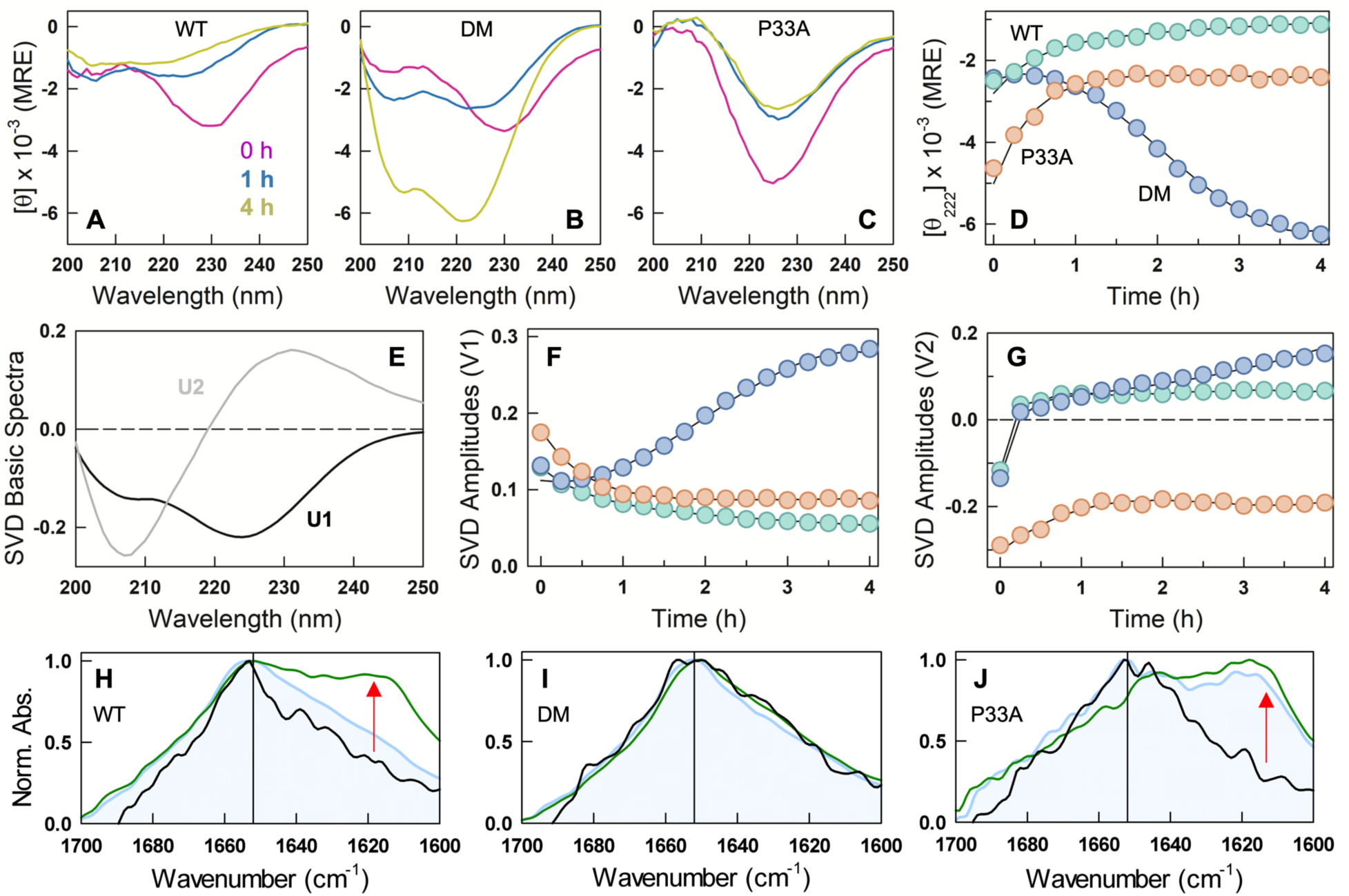
Structural changes in polyP-induced assemblies. (A-C) Far-UV CD spectra of 55 µM WT (A), DM (B), and P33A (C) at different time points - 0 hour (pink), 1 hour (blue), and 4 hours (yellow) - after the addition of 22 µM polyP at 298 K and displayed in mean residue ellipticity (MRE) units of deg. cm^2^ dmol^−1^. (D) Time dependence of the signal at 222 nm for the variants studied. Note the clear secondary-structure acquisition in the DM with time. (E) Basis spectra from a global singular value decomposition (SVD) of time-wavelength far-UV CD data. The first and second components are shown in black and gray, respectively. (F, G) Amplitudes of first (F) and second (G) spectra as a function of time for CytR variants following the color code in panel D. (H-J) FTIR spectra for WT (H), DM (I), and P33A (J) showing normalized absorbance recorded at the wavenumber range of 1700 – 1600 cm^−1^. Black curves represent the FTIR spectra of protein in the absence of polyP. The blue and green curves are the FTIR spectra of protein after the addition of polyP at 0 min and 60 min, respectively. Red arrows indicate the change in peak intensity after the addition of polyP, showing beta-sheet-like conformation in WT and P33A between 1610-1630 cm^−1^. The vertical black line is at 1652 cm^−1^, the amide I frequency indicative of helical structure.

Since the spectra measured with polyP are an effective average of structural signatures from both the condensed- and dilute-phases and of different secondary-structure types, the spectral shapes cannot be directly interpreted. To extract structural signatures and the associated trends, we perform a global singular value decomposition (SVD) of the spectral time series of the three variants. The first two components contribute 92% of the overall signal change (Figure 4E). Of this, the first component is the spectrum of an *⍺*-helix with clear minimum at 222 nm and another at 208 nm, and accounts for 71% of the signal change. The amplitude of this component decreases with time in the WT and P33A variant, implying that helical conformations are less favored in their assemblies; this feature also signals the population of an alternate structural form which is observable in the second component (Figure 4F). In the DM, the amplitude of the helical component increases with time as expected when the droplet dissolves due to preferential transport into the dilute phase. However, the helical content recovers to just ∼15% at 4 hours (Figure 4F), conceivably due to non-specific interaction with polyP resulting in partially structured states or due to a small fraction of oligomers.

The second component, accounting for 21% of the signal change, bears the features of polyproline II structures in unfolded conformations^51^ with a maximum at 231 nm and a minimum at 207 nm (Figure 4E). There is a marginal increase in the amplitude of this component in the WT in the first hour following which it stabilizes (Figure 4G). The amplitude of this component increases in the DM continuously even at 4 hours indicating a tussle between the helical and polyproline II-like structures (Figure 4G). Since the droplets are fully dissolved at 4 hours and by which time the ThT fluorescence starts to marginally increase, it is likely that smaller-sized aggregates (as the contribution of this component is only ∼20% of the overall change) coexist with folded, and partially structured helical conformations in the DM-polyP mixtures at longer times. In the P33A variant, the amplitude of second component has a negative sign and when multiplied with the second component (required for reconstructing the original spectra) points to a strong contribution from *β*-turn or polyproline I conformations, mixed with *β*-sheet.^52^ This is further confirmed by FTIR spectroscopy on P33A wherein a strong absorbance band is evident in the wavenumbers 1610 – 1630 cm^−1^, which is absent in both the WT and DM at *t* = 0 (Figure 4H-4J). At *t* = 60 minutes, the relative intensity of this band increases in the WT and P33A, while no change in spectral shape is evident for the DM.

In summary, we find that both the WT and the DM ‘unfold’ on forming condensates with polyP, with polyproline II-rich structures observed at longer times (∼1-2 hours) in both the mixtures. The condensates thus formed are metastable, either undergoing a liquid-to-solid transition as in the WT or partially dissolving with time as in the DM. At the 4-hour time point, however, the DM acquires helical character due to the partial dissolution of the condensates while the WT forms only aggregates. The starting ensemble thus determines the extent of metastability and the maturation of condensates, and not just the nature of assemblies formed.

### DNA is insensitive to the starting conformational ensembles

DNA is the primary anionic polymer bound by CytR, albeit without sequence preference and involving a distribution of binding free energies.^41^ We find that CytR WT also forms condensates with just 1 *μ*M of double-stranded DNA in a concentration-dependent manner at 20 mM phosphate buffer, pH 7 (Figure 5A, 5B). The droplets are liquid-like displaying coalescence behavior (Figure 5C), recovering fully on photobleaching in a few seconds and with a FRAP *t*_1/2_ of 5.5 ± 0.7 seconds (green in Figure 5D, 5E), comparable to the droplets formed in the presence of polyP. Both the DM and the P33A variants form phase-separated condensates with DNA (Figure 5F, 5G), unlike aggregated assemblies observed with polyP-P33A droplets (Figure 2G).

**Figure 5.**
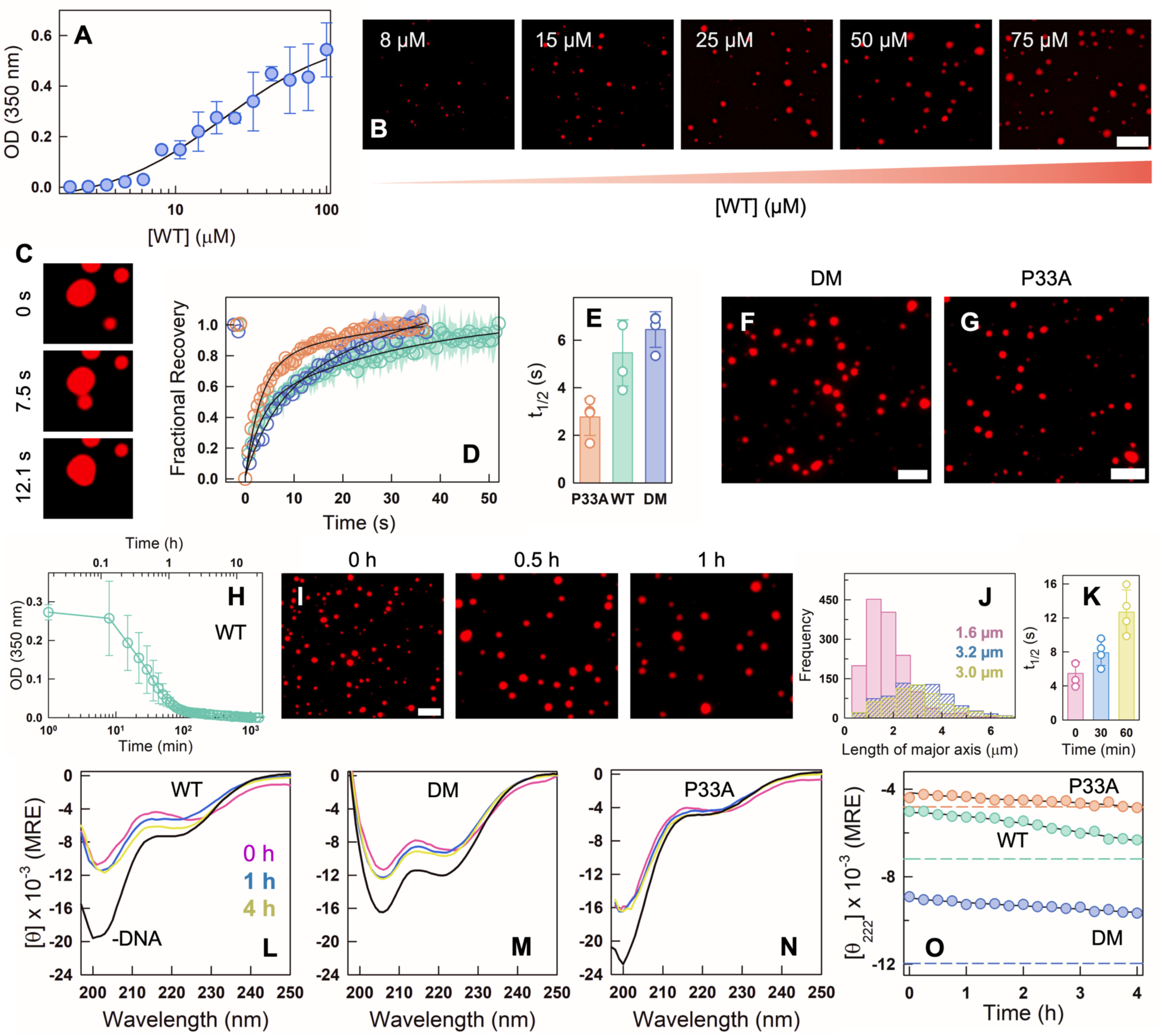
DNA induces metastable condensates that dissolve with time. (A) Changes in turbidity in solutions containing increasing concentrations of the WT and a fixed 1 µM concentration 45-bp specific DNA. The data is the mean from N=2 experiments, and the error bar represents the spread. (B) Fluorescence microscopy images of NHS-rhodamine labeled WT at different concentrations with 1 µM DNA. The scale bar is 10 µm. (C) Representative time-lapse images of WT showing a fusion event. (D) FRAP recovery curves of NHS-rhodamine-labeled CytR WT (green), DM (blue), and P33A (red). The data represents average from N=4 experiments. The experimental errors are shown as shaded areas. (E) Recovery half-times for the CytR variants at the earliest time points (0 minutes). P33A shows the fastest recovery, followed by WT and DM. (F, G) Fluorescence microscopy images of 25 µM DM (F) and P33A (G) in the presence of 1 µM 45 bp DNA. The scale bar is 10 µm. (H) Turbidity of 25 µM WT with 1 µM DNA as a function of time. Data (circles) are from N = 2 experiments and the error bar indicates the spread. (I) Fluorescence microscopy images of NHS-rhodamine labeled WT with DNA at different time points. The scale bar is 10 µm. (J) Droplet size distribution curve for WT in the presence of DNA at different time points - 0 min (pink), 30 min (blue), and 60 min (yellow). Numbers within the plot represent the mean droplet dimensions at the corresponding time points. (K) Recovery half-times from FRAP experiment on WT with DNA at different time points for N=4 experiments following the color code for panel J. (L-N) Far-UV CD spectra of 25 µM WT (left), DM (middle), and P33A (right) at different time points - 0 hour (pink), 1 hour (blue), and 4 hours (yellow) - following the addition of 1 µM 45 bp DNA at 298 K. Black curve is the protein spectra in the absence of DNA. The data is reported in mean residue ellipticity (MRE) units of deg. cm^2^ dmol^−1^. (O) Time-dependent changes in far-UV CD signal at 222 nm for the CytR WT (green), DM (blue), and P33A (red). Dashed lines show MRE signal at 222 nm for the corresponding proteins in absence of DNA.

The time-dependent properties of the condensates with DNA again differ from those of polyP: the condensates dissolve within 90 minutes in both the WT and DM (Figure 5H, 5I), while taking nearly 3 hours for the P33A variant (Figure S3). Numerous droplets with a mean size of 1.6 microns form within the first few minutes (Figure 5J), which coalesce to form larger droplets of size 2-3 microns, following which they dissolve. The FRAP *t*_1/2_ increases by a factor of two at 60 minutes compared to the initial time point (Figure 5K), and a very slow phase is also observed at 60 minutes (Figure S4A), indicating ∼20% immobile fraction. We find that OD is less sensitive to droplet formation with DNA, as we do observe few droplets even at 3 hours, a timepoint at which the OD is effectively zero (Figure S3). Both the DM and P33A mutants recover fully (amplitudes ∼1) and with similar half-times compared to the WT at 60 minutes (Figure S4). To rule out the possibility of concentration differences (22 µM for polyP and 1 µM for DNA) contributing to the observed diverse behavior across the two anionic polymers, we carried out a set of control experiments at 11.2 µM DNA, a concentration at which the number of negative charges are equivalent to that of 22 µM polyP. However, the trends were identical to that observed at lower DNA concentration (Figure S5), further underscoring that the differences in chemical and conformational properties of the anionic polymers contribute to the diverse assembly behaviors.

None of the three proteins ‘unfold’ in the DNA-driven condensates; there is a marginal reduction (i.e. the signal becomes less negative) in the far-UV CD signal at *t* = 0 following which it quickly recovers to reach a value near that of the CD signal in the absence of DNA (Figure 5L-5M). The absence of large structural change or ageing into an aggregate with time is also evident in FTIR spectra which do not change significantly (Figure S6). The WT, however, does display 50% less negative signal at ∼202-204 nm which shifts to longer wavelength after the first hour (blue and green spectra in Figure 5L), indicating helical structure acquisition and mirroring earlier reports.^36,37^ At higher DNA concentrations as used in the current study, this structural shift appears to be offset by unfolding in other regions likely through non-specific interactions. This leads to little signal change at 222 nm – note that this shift is not observable in the two mutants – and thus requiring detailed atomic-level studies to discern the subtle structural modulations induced by DNA at longer times.

To further test the lack of sensitivity of DNA to the starting conformational ensemble, we carried out a similar time-dependent experiment with polyP and DNA, but with the folded DBD of FruR (fructose repressor; Figure 6A) and the molten-globular variant Y19A FruR.^53^ PolyP spontaneously induces aggregation in FruR, while the molten-globular variant forms condensates but only at the earliest times (Figure 6B, 6C, 6D). The intensity of ThT fluorescence protein-polyP mixture increases with time indicating that polyP induces a time-dependent enhanced aggregation of FruR, while the FruR Y19A-polyP mixture undergoes a rapid liquid-to-solid transition within the first 20 minutes (Figure S7). As with the CytR WT, the change in turbidity on aggregation happens earlier than the increase in ThT fluorescence (Figure 6C). The far-UV CD spectral patterns are different in the presence and absence of polyP, with the WT displaying altered secondary structure signature in the presence of polyP (Figure 6E, 6F). The Y19A mutant shows a significantly weakened helical structure content signaling structural destabilization and hence unfolding within the condensate (Figure 6G, 6H). In the presence of DNA, however, both the folded and the molten-globular variants display no change in turbidity with time (Figure 6I – 6L).

**Figure 6.**
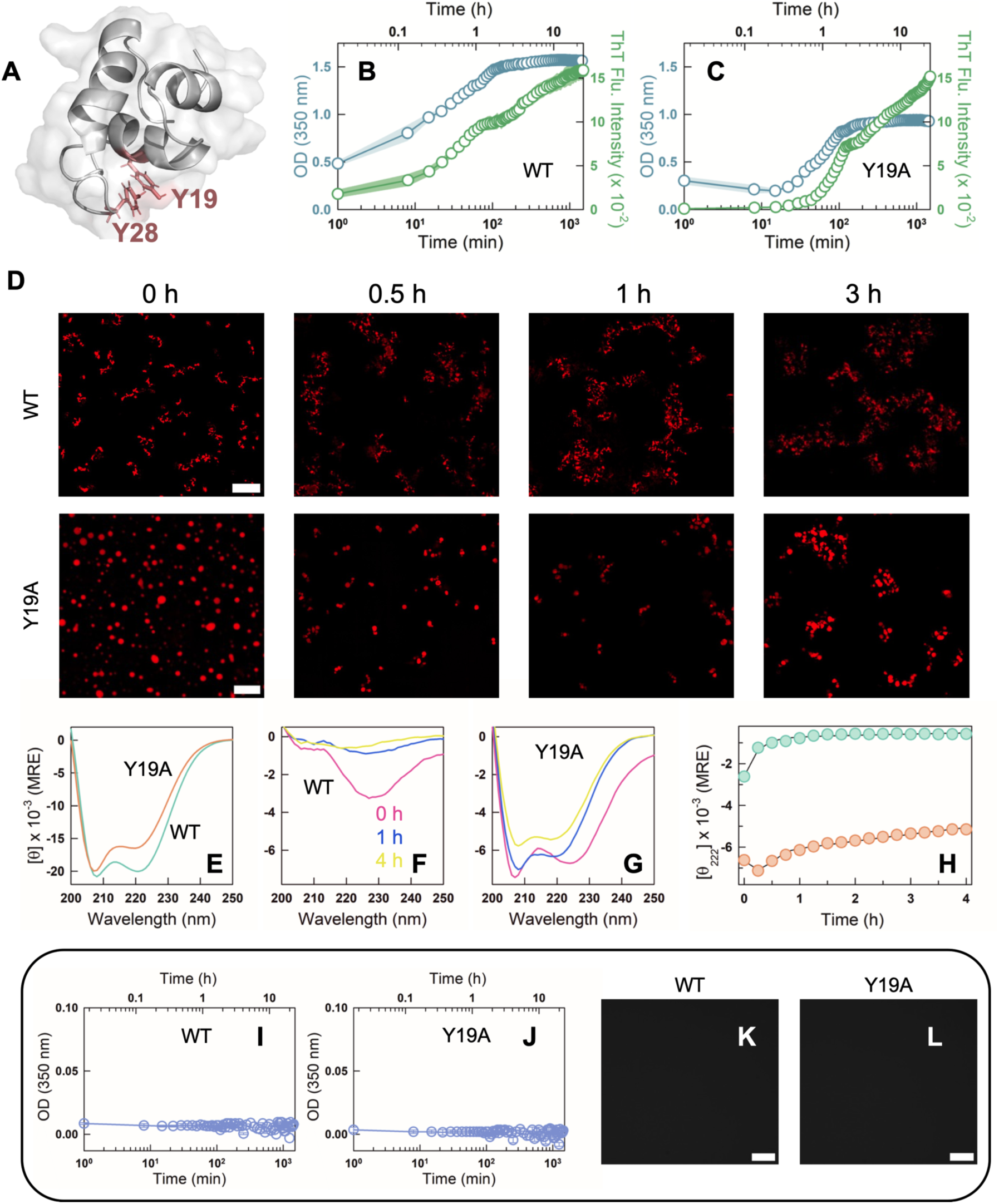
PolyP is able to discriminate between folded and molten-globular variants of FruR DBD while DNA does not. (A) 3D structure of the DNA binding domain of FruR highlighting an aromatic stacking interaction between Y19 and Y28. (B, C) Turbidity (blue circles and left y-axis) and ThT fluorescence intensity (green circles and right y-axis) curve for 90 µM FruR WT (B) and Y19A (C) in the presence of 22 µM polyP. The mean data from N= 2 experiments and the corresponding spread are shown as circles and shaded area, respectively. (D) Representative fluorescence microscopy images of NHS-rhodamine labeled FruR WT (top panel) and Y19A (bottom panel) in the presence of polyP at different time points (0, 0.5,1 and 3 hours). The scale bar is 10 µm. (E) Far-UV CD spectra of FruR WT (green) and Y19A (red) at 298 K in mean residue ellipticity (MRE) units of deg. cm^2^ dmol^−1^ in the absence of polyP. (F and G) Far-UV CD spectra of 55 µM FruR WT (F) and Y19A (G) at different time points - 0 hour (pink), 1 hour (blue), and 4 hours (yellow) - after the addition of 22 µM polyP at 298 K and displayed in mean residue ellipticity (MRE). Note that the y-axis scales of this panel is different from panel E. (H) Time dependence of the signal at 222 nm for WT (green) and Y19A (red). (I and J) Turbidity of 25 µM WT (I) and Y19A (J) with 1 µM DNA as a function of time from N = 2 experiments. No changes in turbidity are observed with time. (K and L) Representative fluorescence microscopy images of NHS-rhodamine labeled FruR WT (K) and Y19A (L) in the presence of DNA. The scale bar is 10 µm.

## Conclusions

Phase separation of disordered CytR with the anionic polyphosphate is quite robust to solution conditions. However, when the native ensemble is perturbed via mutations to result in more order (DM) or less order (P33A) relative to the WT, polyP induces either phase separation or aggregation, respectively (Figure 2). The time-dependent behavior is markedly diverse with liquid-to-solid transition, partial dissolution and enhanced aggregation in CytR, DM and P33A, respectively (Figure 3). The trends are consistent across the different experimental protocols employed including turbidity/fluorescence experiments, fluorescence microscopy, far-UV CD and FTIR (Figures 2 - 4). On the other hand, DNA forms only metastable condensates that dissolve with time, despite different degrees of structural order in the native ensemble of CytR variants (Figure 5). To further prove that polyP interacts differently with ensembles, we performed experiments on the molten-globular variant of FruR (Y19A mutant) and the folded wild-type protein; we see two different behaviors with the WT aggregating and the mutant undergoing a liquid-to-solid transition with polyP, while DNA does not form either aggregates or droplets (Figure 6).

The driving force for the different assembly processes is expected to have a strong contribution from the strength of intermolecular charge-charge interactions, the relative entropic penalty for inducing a conformational change in polyP versus that in the protein, release of counterions and restructuring of water. The balance between these different enthalpic-entropic terms results in condensates being more kinetically accessible phase but which are inherently metastable (as in the CytR WT, DM and FruR Y19A), as a strong time dependence is visibly observable even in the first few minutes. With time, they access the globally stable phase which is either the partially dissolved condensate or the more aggregated phase, conditions at which the chemical potential of the different constituent molecules equalize across the phases. Detailed molecular simulations that capture the relative experimental trends observed across the mutants are required to extract the contributions of each of the energetic and entropic terms.

Our observations establish that polyP is more sensitive to the conformational features of proteins than DNA, thereby contributing to the diverse outcomes. PolyP is expected to be highly flexible owing to its relatively simple chemical structure with only phosphoanhydride bonds that link adjacent units which are highly polar, while DNA has bulkier sugar and the apolar bases that limit its flexibility. It is therefore possible that polyP, owing to its flexibility, is able to wrap around the protein surface or even induce structural changes in proteins in a distinct manner depending on the nature of the starting ensemble. Strong evidence for the latter comes from our far-UV CD studies wherein we find that the helical signature of the CytR variants is fully lost on adding polyP (Figure 4). Our results are consistent with previous observations of structural changes and ‘unfolding’ of proteins within condensates.^54–57^ We additionally show that polyproline-II-like conformations are more probable within condensates formed by the CytR WT and the DM (Figure 4). One other observation stands out - the ThT fluorescence increases upon aggregation at a later time point than the changes in OD in CytR WT/DM and FruR Y19A; this is evidence that smaller aggregates are better captured by OD changes than ThT fluorescence, as the latter is more sensitive to larger aggregates.

The large charge density associated with polyphosphates and their natural abundance makes them prime candidates for inducing phase separation^58^ through complex coacervation with proteins rich in basic amino acids. In fact, most DBDs which are typically associated with transcription factors are rich in basic amino acids. This opens up a fascinating possibility where polyP and DNA could compete with transcription factors or work together forming ternary condensates regulating gene expression, as shown in studies on the bacterial protein Hfq.^27^ Taken together with our results on CytR, caution needs to be exercised in interpreting in-cell microscopy results, as it is possible that polyP is involved in condensates, especially when proteins of interest are rich in basic residues. Finally, the full length CytR plays an important role in transcription regulation of genes involved in stress response, including critical roles in nucleoside catabolism, context-dependent activation or repression, and pathogenesis.^59–61^ CytR also binds to the promotor region of the rpoH gene and represses transcription, thus regulating the expression of σ^32^ – an RNA polymerase subunit essential for the transcription of heat shock proteins.^62,63^ Given the results of the current work and since polyP concentrations rise significantly in stressed cells,^20^ it is tempting to speculate that polyP interacts with CytR to form condensates and sequesters it, thus promoting σ^32^ transcription. Studies on CytR combining other stress response regulators, specifically with the nucleoid-associated sensory protein H-NS, with DNA and polyP could unravel hitherto unexplored mechanisms regulating the nuanced gene expression patterns in enterobacteria.

## Abbreviations

polyP: polyphosphate
CytR: cytidine repressor
LLPS: liquid-liquid phase separation
DIC: differential interference contrast
CD: circular dichroism
FTIR: Fourier-transform infra-red
DNA: deoxy ribonucleic acid

## Acknowledgements

The authors are grateful for the support from the Department of Biotechnology (DBT, India) for the grant BT/PR41973/BRB/10/1967/2021 to A. N. N. The authors acknowledge the FIST facility (SR/FST/LS-II/2020/552(C)) sponsored by the Department of Science and Technology (DST, India) and the ICSR - Common Instruments Facility (CIF) at the Department of Biotechnology, IIT Madras (Chennai, India) for the instrumentation. The authors thank Vijay Rangachari and Ethayaraja Mani for the discussions, Swathi Sudhakar for the help in flow chamber preparation, and Thalappil Pradeep and Sonali Seth for help in FTIR experiments. D.R. acknowledges Women Leading IITM for the funding.

