## Supporting Information for "Polyphosphate Discriminates Protein Conformational Ensembles More Efficiently than DNA Promoting Diverse Assembly and Maturation Behaviors"

AUTHOR INFORMATION

**Corresponding Authors**

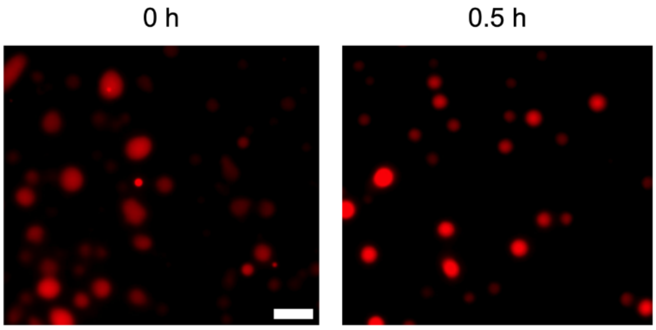


**Figure S1** Fluorescence microscopy images of 90 µM WT with 22 µM polyP at 160 mM ionic strength (pH 7) immediately after adding polyP (left panel) and after 30 minutes (right panel). The scale bar is 10 µm.

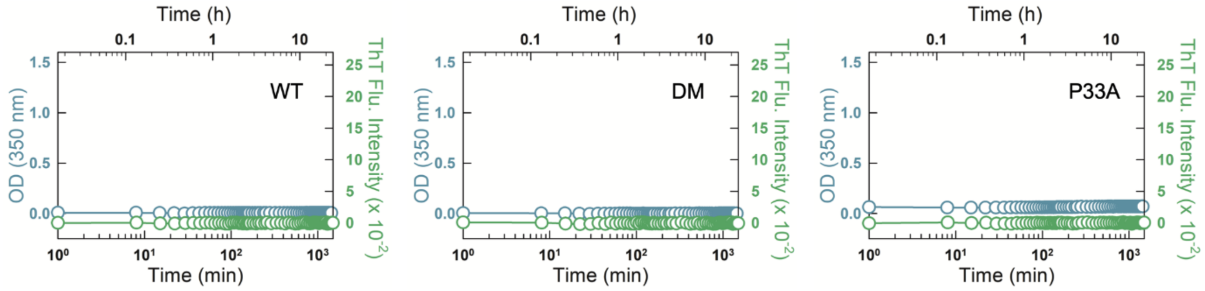


**Figure S2** Turbidity (blue circles and left y-axis) and ThT fluorescence intensity (green circles and right y-axis) curve for 90 μM WT (left panel), DM (middle panel), and P33A (right panel) in the presence of 5 μM ThT.

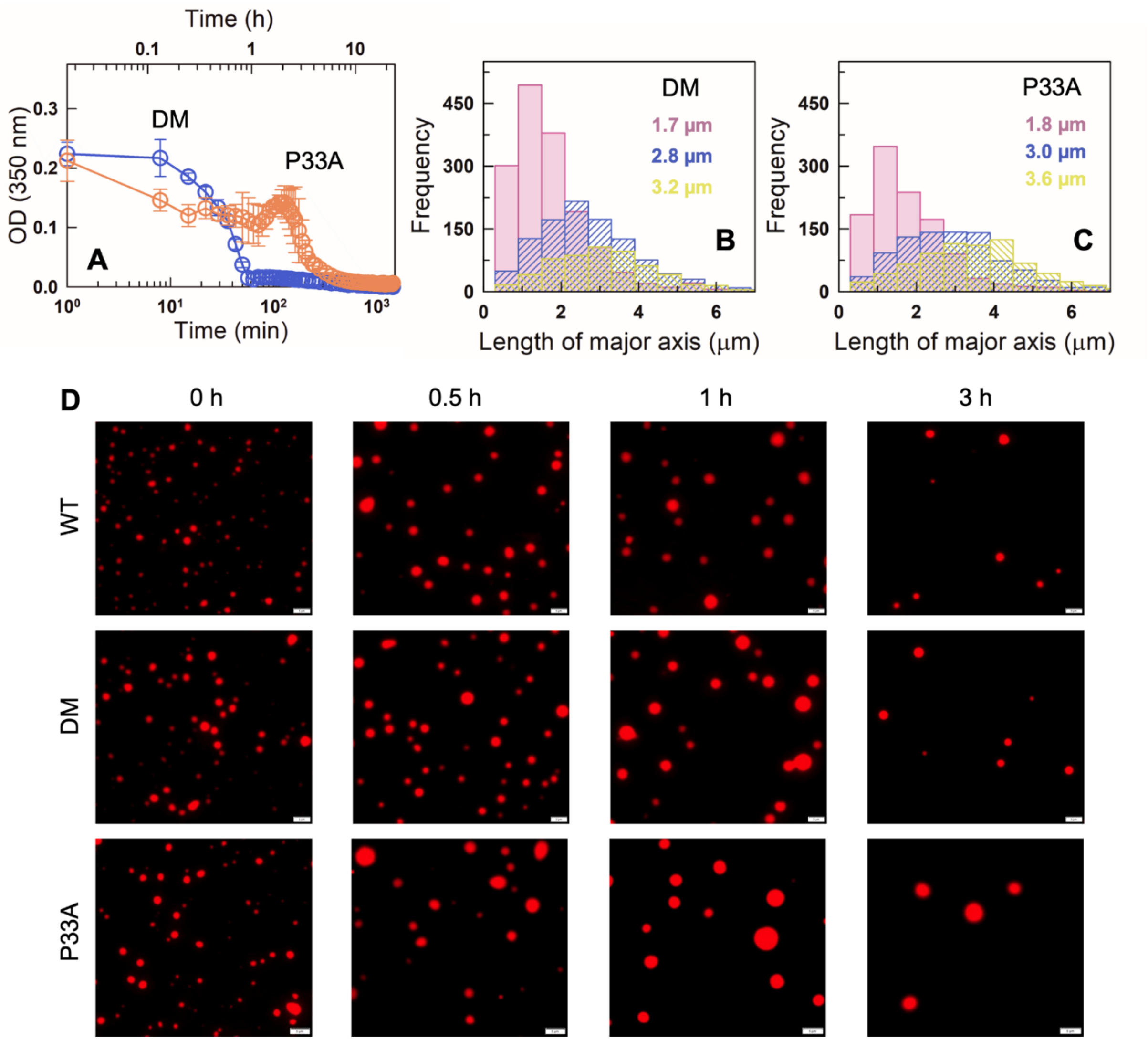


**Figure S3** (A) Turbidity of 25 µM DM (blue) and P33A (orange) with 1 µM DNA as a function of time. The mean and the spread from N = 2 experiments are shown as circles and error bar, respectively. (B, C) Droplet size-distributions for DM (B) and P33A (C) in the presence of DNA at different time points: 0 min (pink), 30 min (blue), and 60 min (yellow). Values given in the plot represent the mean dimensions of droplets at the corresponding time points. (D) Fluorescence microscopy images of NHS-rhodamine labeled WT (top), DM (middle) and P33A (bottom) in the presence of 45 bp DNA at different time points. The scale bar is 5 µm.

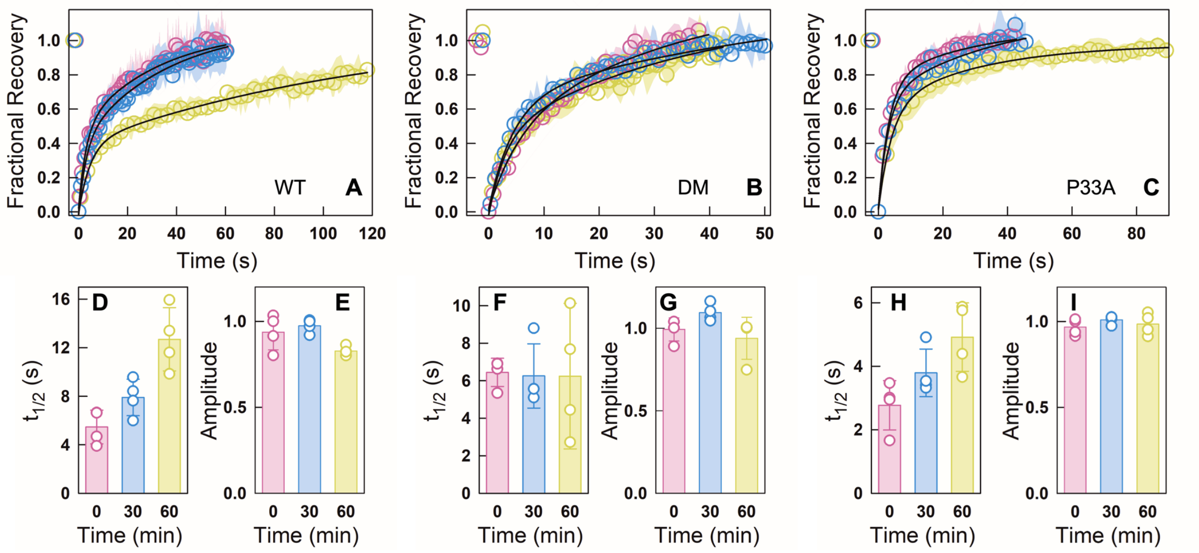


**Figure S4** FRAP experiments for CytR variants in the presence of DNA at different time points: 0 min (pink), 30 min (blue), and 60 min (yellow). Data are reported as mean ± s.d. for N = 4 experiments. (A-C) FRAP recovery curves of NHS-rhodamine labeled WT (A), DM (B) and P33A (C). The experimental errors are smaller than the size of data points. (D, F, H) FRAP recovery half-times for WT (D), DM (F), and P33A (H) at the indicated time points. (E, G, H) Recovery amplitude plots for WT (E), DM (G) and P33A (I) at different time points.

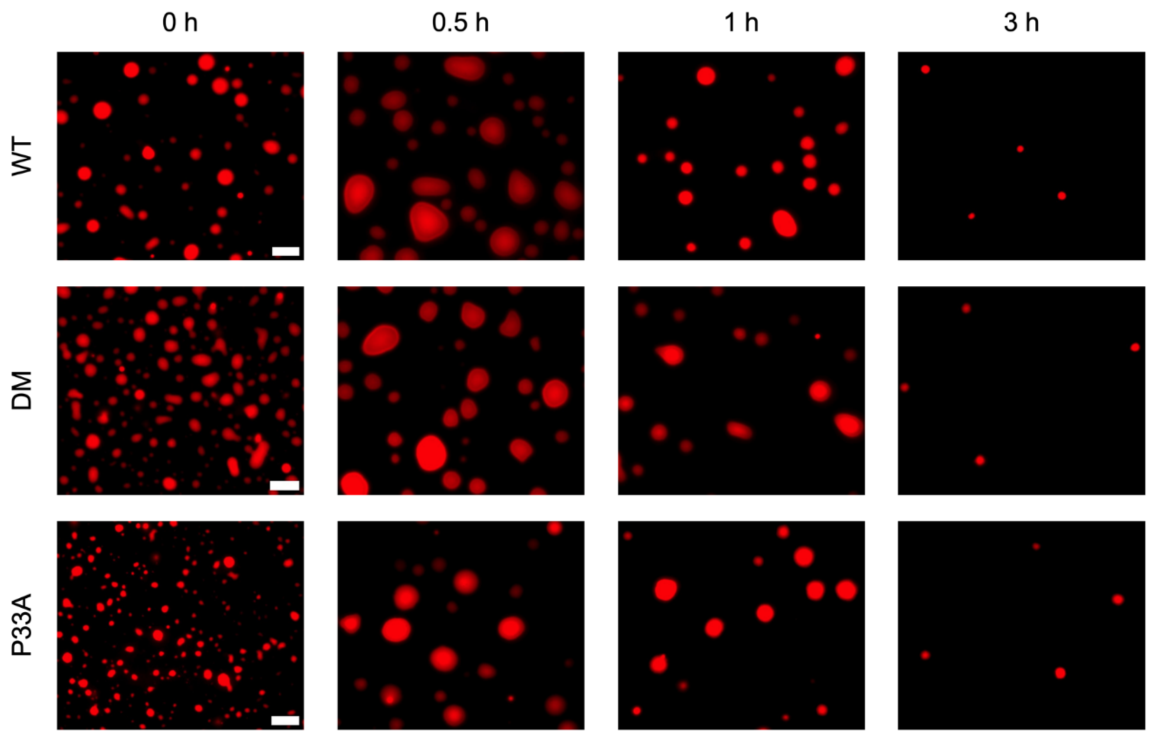


**Figure S5** Fluorescence microscopy images of 90 µM NHS-rhodamine labeled WT (top), DM (middle) and P33A (bottom) in the presence of 11.24 µM of 45-bp DNA at the time points indicated. The scale bar is 10 µm.

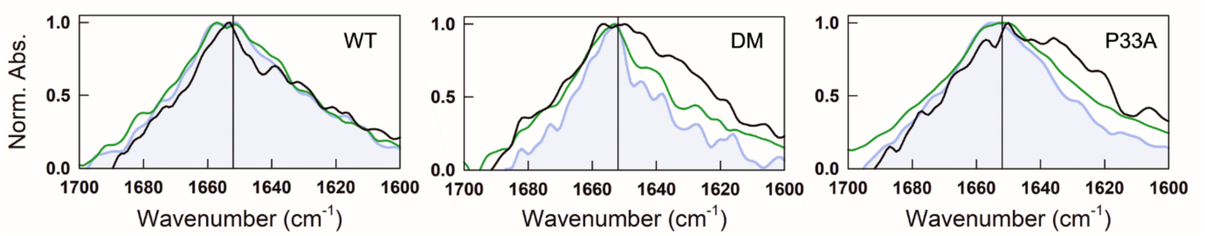


**Figure S6** FTIR spectra of WT (left), DM (middle), and P33A (right) showing normalized absorbance recorded at a wavenumber range of 1700 – 1600 $cm^{-1}$. Black curves represent FTIR spectra of protein in the absence of DNA, and blue and green curves show FTIR spectra of protein after the addition of DNA at 0 min and 60 min, respectively.

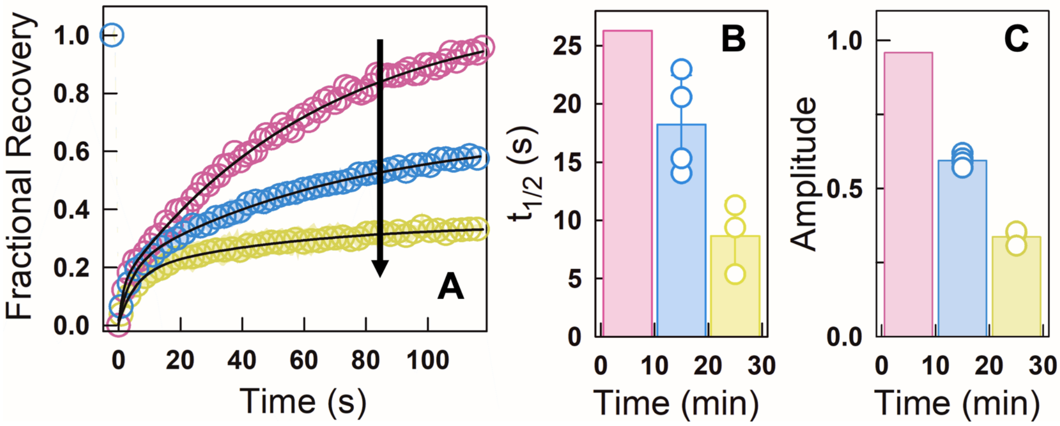


**Figure S7** FRAP experiments on Y19A variant of FruR in the presence of polyP at different time points: 0 - 10 min (pink), 10-20 min (blue), and 20-30 min (yellow). (A) FRAP recovery curves of NHS-rhodamine labeled FruR. Arrow indicates the movement of the recovery profiles at longer incubation times. (B) Recovery half-time at the indicated time points. (C) Recovery amplitude at different time points. FRAP data for 0-10 min represents data from a single experiment as the condensates of Y19A are highly metastable (also see Figure 6C). The data acquired between 10-20 and 20-30 minutes are shown as mean ± s.d from N = 4 and 3 experiments, respectively. The experimental errors are smaller than the size of data points.
